# A new contact killing toxin permeabilizes cells and belongs to a large protein family

**DOI:** 10.1101/2021.03.31.437997

**Authors:** Cristian V. Crisan, Harshini Chandrashekar, Catherine Everly, Gabi Steinbach, Shannon E. Hill, Peter J. Yunker, Raquel R. Lieberman, Brian K. Hammer

## Abstract

*Vibrio cholerae* is an aquatic Gram-negative bacterium that causes severe diarrheal cholera disease when ingested by humans. To eliminate competitor cells in both the external environment and inside hosts, *V. cholerae* uses the Type VI Secretion System (T6SS). The T6SS is a macromolecular weapon employed by many Gram-negative bacteria to deliver cytotoxic proteins into adjacent cells. In addition to canonical T6SS gene clusters encoded by all sequenced *V. cholerae* isolates, strain BGT49 encodes an additional locus, which we named auxiliary cluster 4 (Aux 4). The Aux 4 cluster is located on a mobile genetic element and can be used by killer cells to eliminate both *V. cholerae* and *Escherichia coli* cells in a T6SS-dependent manner. A putative toxin encoded in the cluster, which we name TpeV (*<u>T</u>ype VI <u>P</u>ermeabilizing <u>E</u>ffector <u>V</u>ibrio*), shares no homology to known proteins and does not contain motifs or domains indicative of function. Ectopic expression of TpeV in the periplasm of *E. coli* permeabilizes cells and disrupts the membrane potential. Using confocal microscopy, we confirm that susceptible target cells become permeabilized when competed with killer cells harboring the Aux 4 cluster. We also determine that *tpiV*, the gene located immediately downstream of *tpeV*, encodes an immunity protein that neutralizes the toxicity of TpeV. Finally, we show that TpeV homologs are broadly distributed across important animal and plant pathogens and are localized in proximity to other T6SS genes. Our results suggest that TpeV is a toxin that belongs to a large family of T6SS proteins.

**IMPORTANCE:** Bacteria live in polymicrobial communities where competition for resources and space is essential for survival. Proteobacteria use the T6SS to eliminate neighboring cells and cause disease. However, the mechanisms by which many T6SS toxins kill or inhibit susceptible target cells are poorly understood. The sequence of the TpeV toxin we describe here is unlike any previously described protein. We demonstrate that it has antimicrobial activity by permeabilizing cells, eliminating membrane potentials and causing severe cytotoxicity. TpeV homologs are found near known T6SS genes in human, animal and plant bacterial pathogens, indicating that the toxin is a representative member of a broadly distributed protein family. We propose that TpeV-like toxins contribute to the fitness and pathogenicity of many bacteria. Finally, since antibiotic resistance is a critical global health threat, the discovery of new antimicrobial mechanisms could lead to the development of new treatments against resistant strains.

## INTRODUCTION

The Type VI Secretion System (T6SS) is a common bacterial weapon employed by many killer Gram-negative bacteria to translocate toxic protein effectors into adjacent target cells (1, 2). The harpoon-like proteinaceous apparatus is anchored to the membrane of killer cells by the membrane complex, which spans the inner membrane and periplasm (3–5). Hcp (hemolysin-coregulated protein) hexamers stack to form an inner tube that is capped at the distal end by a trimer of VgrG (valine-glycine repeat protein G) tip-forming proteins (2, 6, 7). PAAR (proline-alanine-alanine-arginine) proteins also interact with VgrGs and expand the toxin repertoire (8, 9). The T6SS uses a contraction mechanism that propels the inner tube and delivers the toxic payload (10–12).

*Vibrio cholerae* is a wide-spread gastrointestinal pathogen that has caused seven cholera pandemics (13). The bacterium is found in polymicrobial marine ecosystems in association with copepods, fish and insects (14–16). To colonize hosts and survive in environmental settings, *V. cholerae* employs T6SS effectors that disrupt the cell envelope of competitor cells (17–24). T6SS genes are distributed across a large cluster and two auxiliary clusters in all sequenced *V. cholerae* isolates (25, 26). In clinical strains like V52 and C6706, the large gene cluster encodes a VgrG tip-forming protein with a C-terminal peptidoglycan-degrading domain (23). Auxiliary clusters 1 and 2 encode the TseL lipase and VasX colicin-like effectors, respectively (19–22). An auxiliary cluster 3 is found in a subset of *V. cholerae* isolates and contains a peptidoglycan-degrading toxin (27–29).

Although most clinical *V. cholerae* strains encode T6SS effectors with conserved activities, *V. cholerae* strains obtained from sources other than patients harbor a more diverse repertoire of T6SS toxins (25, 26, 30, 31). We previously identified auxiliary 5 (Aux 5) T6SS clusters in several *V. cholerae* strains, which encode predicted phospholipase effectors (25). Recently, several *V. cholerae* strains have been shown to possess an Aux 6 T6SS cluster with antibacterial activity (31). We and others have also reported that many *V. cholerae* strains (but not C6706) contain an additional gene cluster with putative T6SS components, which we named Aux 4 (25, 32). However, the activity of the cluster, the roles played by the encoded genes in microbial competition and the toxicity of the putative effector have not been validated.

Here we demonstrate that the Aux 4 cluster can be used by *V. cholerae* to kill bacterial cells in a T6SS-dependent manner. We report that the toxin found within the cluster permeabilizes cells and disrupts the membrane potentials when expressed in the periplasm of *Escherichia coli* cells. A protein encoded by a gene found immediately downstream of the effector neutralizes its toxicity and acts as a protective immunity factor. Finally, we show that homologs of the Aux 4 effector are found in diverse bacterial species, including human, animal and plant pathogens. The potent antimicrobial activity of TpeV and broad distribution of identified homologs suggest the toxins confer significant competition advantages to bacteria that harbor them.

## RESULTS

### The Aux 4 tpeV-tpiV are an active effector-immunity pair in strain BGT49

*V. cholerae* strain BGT49 encodes the Aux 4 cluster, in addition to the canonical T6SS large operon and auxiliary clusters 1 and 2 (Fig. 1A). The Aux 4 cluster contains predicted T6SS genes: an *hcp*, a *vgrG*, a DUF4123 chaperone, and a *paar* (33) (Fig. 1A). Genes coding for a putative effector toxin (which we name *tpeV, Type VI Permeabilizing Effector Vibrio*, see below) and a putative immunity protein (which we name *tpiV, Type VI Permeabilizing Immunity Vibrio*, see below) are also found within the cluster (Fig. 1A) (32, 34, 35). The *vgrG* gene does not contain a toxic C-terminal domain, as described for the *V. cholerae* VgrG-1 or VgrG-3 (23, 36, 37). Genes for a restriction modification system are found upstream of the Aux 4 cluster (Fig. 1A). Both the Aux 4 T6SS cluster and restriction modification system genes are flanked upstream by a predicted integrase and downstream by a predicted transposase (Fig. 1A). Attachment (att) sites similar to those found in the *Vibrio* pathogenicity island 1 (VPI-1) also flank the region (32, 38). To experimentally determine that the *tpeV* gene encodes a T6SS toxin, we engineered a *ΔtpeVΔtpiV* target BGT49 strain. The *ΔtpeVΔtpiV* target strain was then co-cultured with either wild type BGT49, *ΔtpeV* (BGT49 lacking the TpeV effector) or BGT49 T6SS-killers. The recovery of the *ΔtpeVΔtpiV* target strain was significantly reduced (by approximately 5 orders of magnitude) when co-cultured with wild type BGT49 killer cells compared to the *ΔtpeV* or T6SS-killer strains (Fig. 1B). This result indicates that TpeV is a T6SS effector that is actively used by *V. cholerae* strain BGT49 to eliminate susceptible cells that lack the TpiV immunity protein.

**Figure 1.**
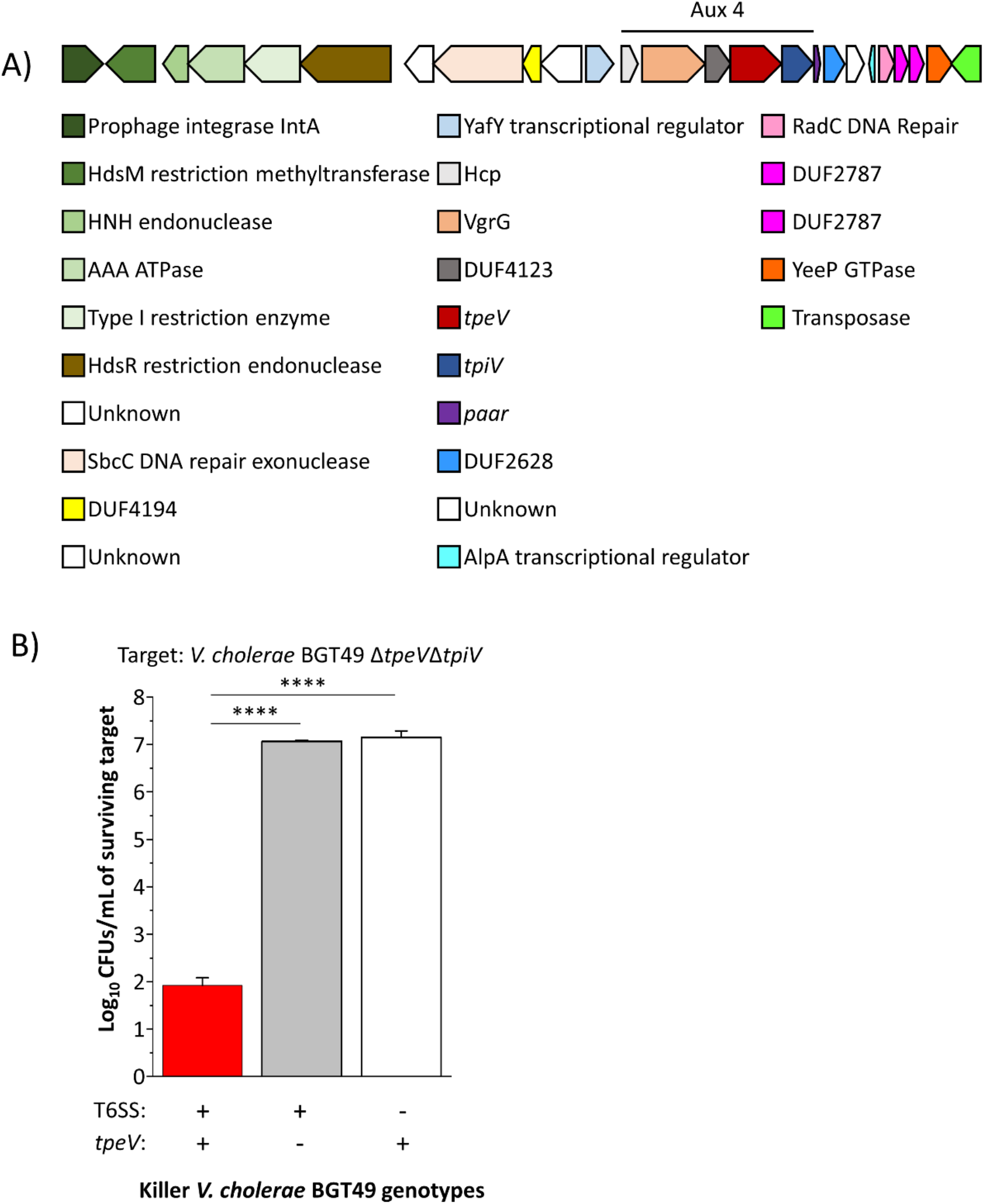
*Vibrio cholerae* strain BGT49 encodes the Aux 4 T6SS cluster and efficiently eliminates target bacteria in a TpeV and T6SS-dependent manner. A) The Aux 4 cluster encodes predicted *hcp, vgrG*, DUF4123-containing chaperone, effector, immunity and *paar* genes. The cluster is found on a predicted mobile genetic element, being flanked by integrase and transposase genes. B) Target *V. cholerae* BGT49 Δ*tpeV*Δ*tpiV* was co-cultured with either WT, Δ*tpeV* or T6SS-killer BGT49. A one-way ANOVA with a post hoc Tukey HSD test was used to determine significance. ****p < 0.0001

### The Aux 4 cluster can be transferred to another V. cholerae strain where it confers competitive advantages

Since we observed that the Aux 4 cluster is located on a predicted mobile genetic element, we hypothesized that it can be used by other *V. cholerae* strains to eliminate competitor cells in a T6SS-dependent manner. In *V. cholerae* C6706, the QstR protein is a gene regulator that is required and sufficient to induce expression of T6SS genes (39, 41). We cloned the Aux4 *vgrG, tap, tpeV, tpiV* and *paar* genes on a plasmid (pAux4) under control of the Ptac promoter. We then introduced the pAux4 plasmid in *V. cholerae* strain C6706*, which constitutively expresses the QstR protein but does not possess Aux 4 cluster genes on its chromosomes (24, 39–41). The *V. cholerae* C6706* killer with the Aux 4 cluster on a plasmid (C6706*/pAux4) efficiently eliminates the parental target strain, unlike a killer C6706* strain carrying a plasmid control (Fig. 2A). By contrast, a C6706*/pAux4 T6SS-strain cannot eliminate the parental target strain (Fig. 2A). To provide further evidence that TpiV can confer immunity, we introduced the *tpiV* gene into target *V. cholerae* C6706 and co-cultured the strain with killer C6706*/pAux4 cells. *V. cholerae* C6706*/pAux4 kills *V. cholerae* target cells with a plasmid control, but not when they encode the *tpiV* gene (Fig. 2B).

**Figure 2.**
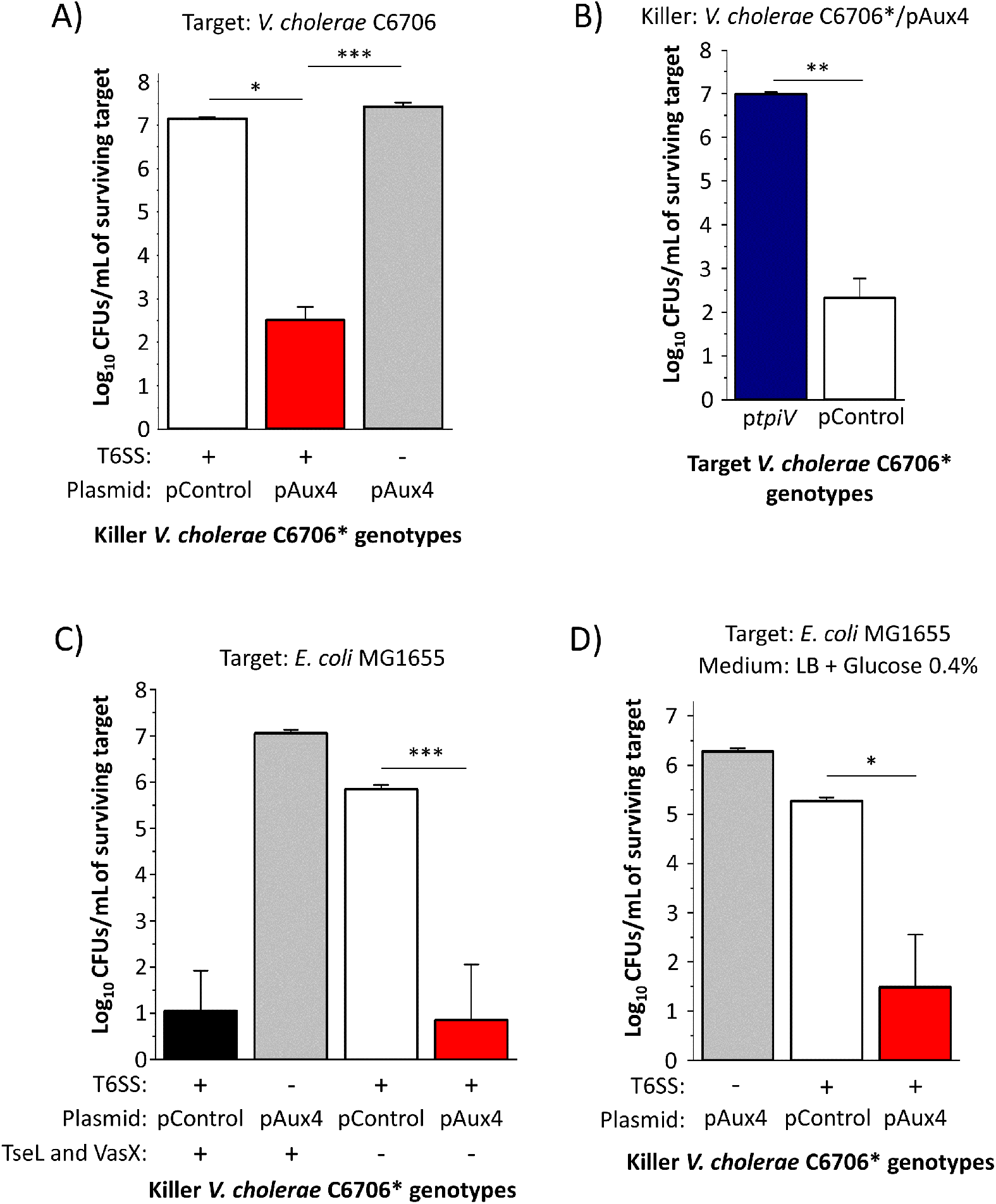
*V. cholerae* C6706* can use the Aux 4 cluster to eliminate target cells in a T6SS-dependent manner. A) *V. cholerae* C6706* (T6SS+ or T6SS-) with a plasmid control or a plasmid encoding the Aux 4 cluster were co-cultured with target parental *V. cholerae* C6706. A one-way ANOVA with a post hoc Tukey HSD test was used to determine significance. B) Killer *V. cholerae* C6706* with the Aux 4 cluster were co-cultured with target C6706 cells with a plasmid control or a plasmid encoding *tpiV*. Welch’s t-test was used to determine significance. C) *V. cholerae* C6706* with deletions in the known *tseL* and *vasX* T6SS effectors containing either a plasmid control or a plasmid with Aux 4 was co-cultured with *E. coli* MG1655 cells. A one-way ANOVA with a post hoc Tukey HSD test was used to determine significance. D) *V. cholerae* C6706* with a plasmid control or a plasmid encoding the Aux 4 cluster were co-cultured with *E. coli* MG1655 on LB medium with 0.4% glucose. A one-way ANOVA with a post hoc Tukey HSD test was used to determine significance. *** p < 0.001, ** p < 0.01 * p < 0.05

We next inquired if the Aux 4 cluster can be used by *V. cholerae* to kill other target bacterial species. A C6706* strain with a functional T6SS that lacks both native TseL and VasX effectors poorly eliminates *E. coli* cells compared to a C6706* strain that harbors both toxins (Fig. 2C) (42, 43). However, the introduction of the Aux 4 cluster into the C6706* strain lacking TseL and VasX effectors restores its ability to efficiently eliminate *E. coli* cells (Fig. 2C). We recently reported that target *E. coli* cells are protected against T6SS attacks from strain C6706* when co-cultured on LB medium supplemented with 0.4% glucose (44). By contrast, we observe that killer C6706*/pAux4 can bypass the glucose-mediated resistance to efficiently eliminate *E. coli* cells even when the co-culture is performed on LB medium with glucose (Fig. 2D). These results confirm that the Aux 4 intoxicates competitor bacterial cells.

### TpeV permeabilizes cells and disrupts the membrane potential

We next used confocal microscopy to examine co-cultures between fluorescently labelled target *V. cholerae* C6706 cells (shown as cyan) and unlabeled killer C6706*/pAux4 cells (Fig. 3A). To each co-culture we added propidium iodine (PI), a molecule that cannot penetrate cells with intact membranes but exhibits high fluorescence when bound to the DNA of cells with compromised membranes. Fluorescently labelled *V. cholerae* target cells are successfully eliminated when co-cultured with killer C6706*/pAux4 cells but remain viable when killer cells cannot assemble the T6SS apparatus (T6SS-) (Fig. 3A). Furthermore, a robust PI signal (depicted with red) is detectable when target cells are co-cultured with C6706*/pAux4 cells (Fig. 3A).

**Figure 3.**
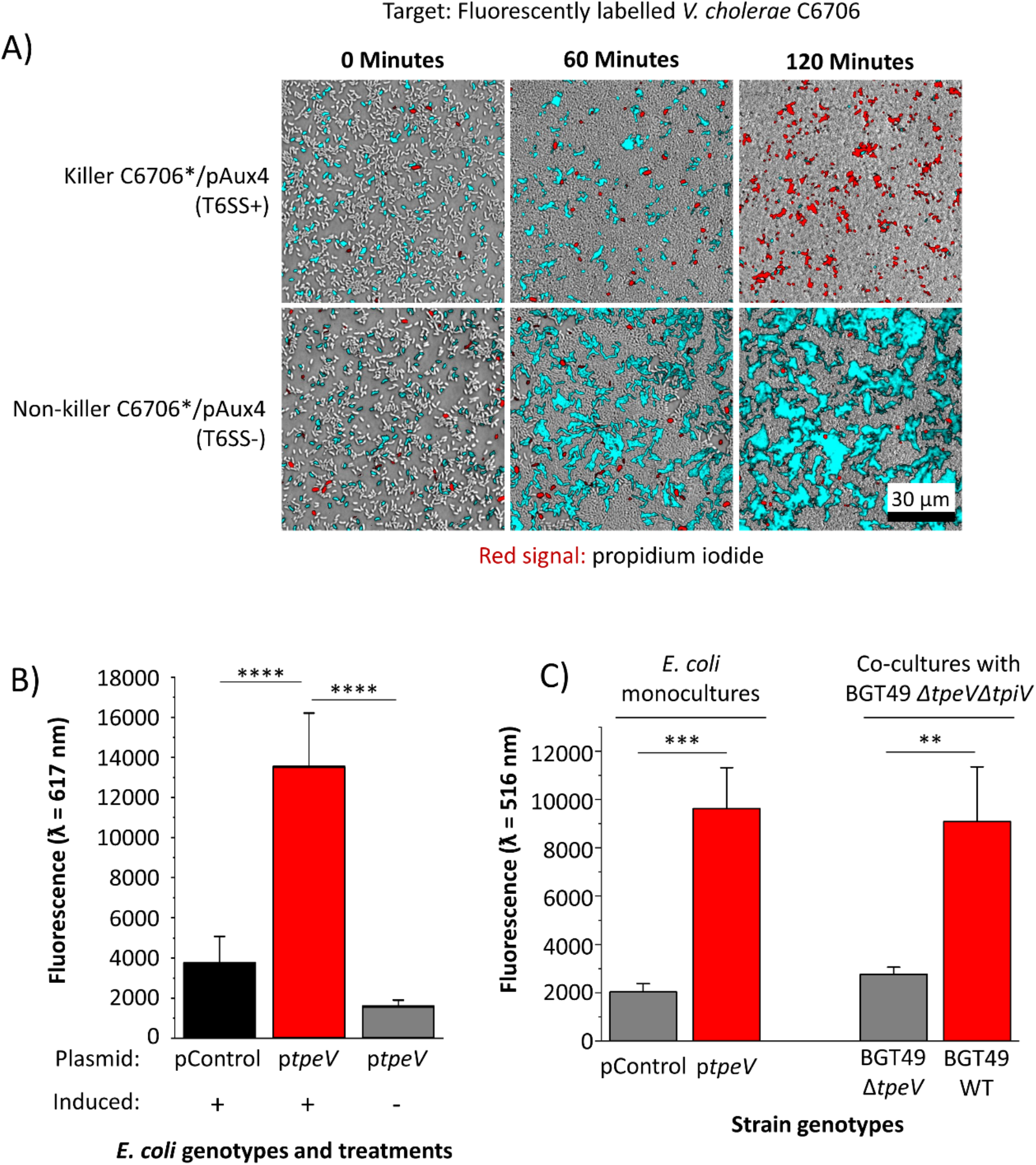
TpeV permeabilizes target cells and disrupts the membrane potential, leading to cytotoxicity. A) Confocal microscopy was used to visualize a co-culture between C6706* cells with Aux 4 (T6SS-or T6SS+) and fluorescently labelled target C6706 cells in the presence of propidium iodide. Scale bar = 30 µm. B) *E. coli* cells carrying a periplasmic *tpeV* construct or plasmid control were incubated with propidium iodide. Fluorescence readings were taken at an excitation λ = 535 nm and emission λ = 617 nm. A one-way ANOVA with a post hoc Tukey HSD test was used to determine significance. C) *E. coli* cells carrying a periplasmic *tpeV* construct or plasmid control, or *V. cholerae* BGT49 co-cultures between target *ΔtpeVΔtpiV* and wild type or *ΔtpeV* killer cells were incubated with the membrane potential-sensitive DiBAC_4_(3) dye. Fluorescence readings were taken at an excitation λ = 490 nm and emission λ = 516 nm. Welch’s t-tests were used to determine significance. **** p < 0.0001, *** p < 0.001, ** p < 0.01

Since we observed that *V. cholerae* cells harboring the Aux 4 cluster can kill and permeabilize target cells in a T6SS-depedent manner, we sought to further characterize the activity of the TpeV effector. The protein does not share primary sequence homology to known toxins and does not contain motifs or domains indicative of function. Tertiary structural prediction algorithms also fail to detect significant homologs with known functions. TpeV has 11 cysteine residues, suggesting that multiple disulfide bonds could play roles in stabilizing the protein. Transmembrane prediction algorithms TMHMM and Phobius do not detect extensive transmembrane regions and SignalP 5.0 does not predict a signal sequence (Supplementary Fig. 1 and 2) (45–47). We also attempted to identify TpeV homologs using the secondary structure predictor Jpred (48). While most homologs are hypothetical proteins with unknown functions, some contain domains similar to the peptidoglycan-binding C-terminal regions of the OmpA protein (49, 50). OmpA proteins are involved in pathogenesis and have diverse functions that include formation of porins and channels (51, 52). Because target *V. cholerae* cells have a substantial PI signal when co-cultured with killer cells harboring the Aux 4 cluster, we hypothesized that TpeV might permeabilize target cells when delivered to the periplasm.

To test this prediction, we introduced plasmid-borne *tpeV* with a periplasmically-directing *pelB* sequence under the control of an inducible promoter into *E. coli* cells. A significantly higher PI signal is detected when *E. coli* cells are induced to express periplasmic TpeV compared to cells that harbor a plasmid control (Fig. 3B). We also hypothesized that TpeV disrupts the bacterial cell membrane potential (19, 53, 54). To test this hypothesis, we used the Bis-(1,3-Dibutylbarbituric acid) Trimethine Oxonol ((DiBAC_4_(3)) potential-sensitive dye, which is excluded from cells with a normal membrane potential but exhibits fluorescence in depolarized cells (54–56). *E. coli* cells that express periplasmically-delivered TpeV have significantly higher DiBAC_4_(3) uptake compared to *E. coli* that express a plasmid control (Fig. 3C). To probe if TpeV can disrupt the membrane potential in a T6SS-dependent manner, we co-cultured *V. cholerae* BGT49 *ΔtpeVΔtpiV* target cells with either BGT49 wild type or BGT49 *ΔtpeV* killers. Following co-cultures with wild type but not BGT49 *ΔtpeV* killers, bacterial membrane potentials are disrupted as cells display an elevated DiBAC_4_(3) signal (Fig. 3C). Taken together, these findings demonstrate that TpeV permeabilizes cells and disrupts the membrane potential of target bacteria.

### TpeV belongs to a large family of T6SS proteins

Since the sequence or predicted structure of TpeV shares no homology to known toxins (including known permeabilizing toxins), we used PHMMER to search if homologs are present in other bacterial species (57). We identified *tpeV*-like genes across diverse Gammaproteobacteria species (Fig. 4, Supplementary Table 1). In all selected bacterial species, genes coding for TpeV homologs are found near known T6SS genes like *hcp, vgrG*, DUF4123-containing chaperones or other structural components (Fig. 4). Our results indicate that TpeV is a representative member of a widely spread family of T6SS toxins with antimicrobial activity that allow cells to eliminate competitor bacteria.

**Figure 4.**
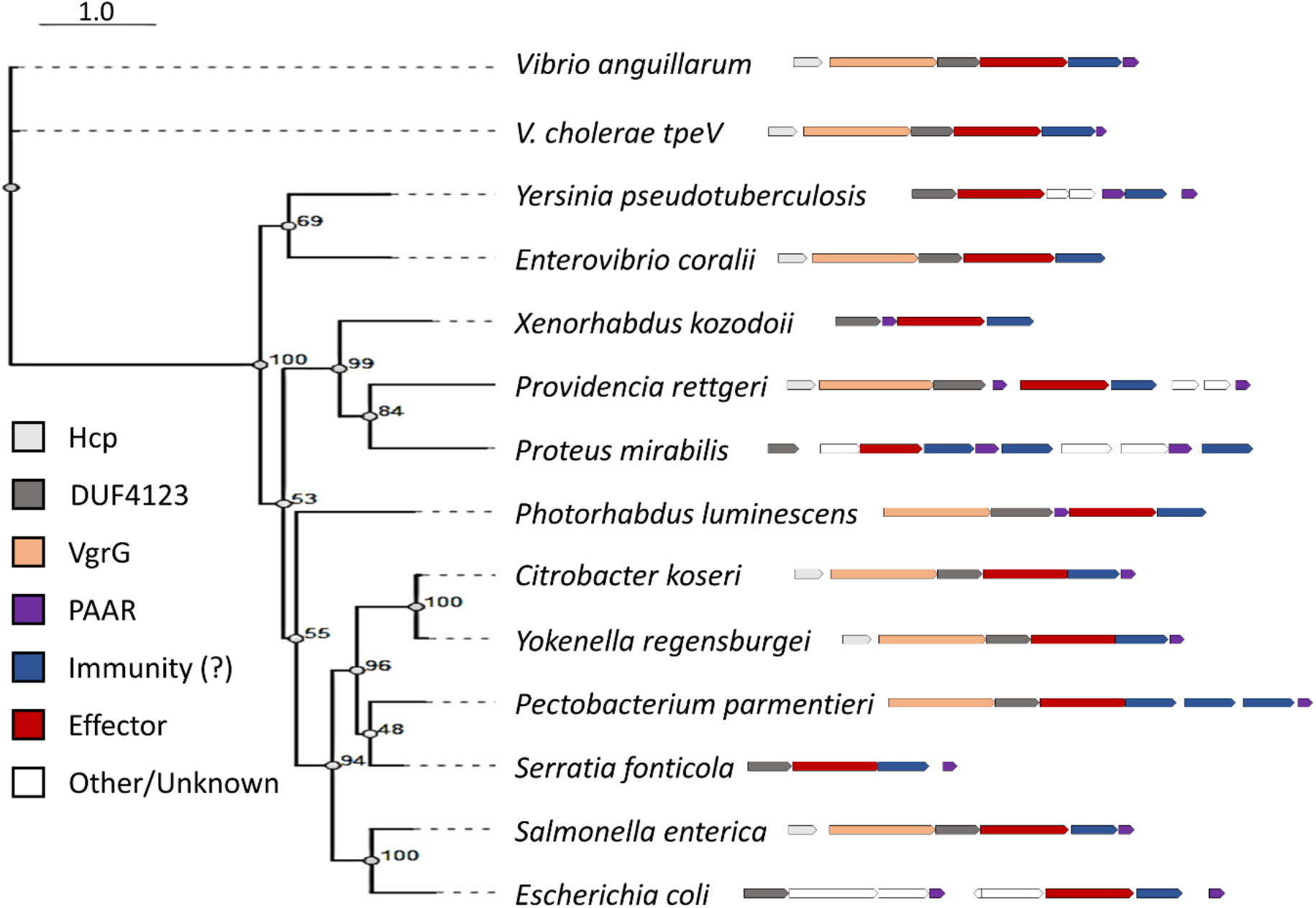
*TpeV* homologs are found in many bacterial species near other T6SS genes. TpeV homologs were identified using PHMMER and selected sequences were aligned using MUSCLE. A phylogenetic tree was constructed with 100 bootstraps.

## DISCUSSION

Here we show that many bacterial species encode homologs of a previously undescribed T6SS protein that intoxicates, permeabilizes and disrupts the membrane potential of target cells. While studies have examined antibacterial effectors from clinical *V. cholerae* isolates such as C6706 and V52, we and others found that strains isolated from sources other than patients encode a more diverse set of putative T6SS toxins (3, 19, 22, 25, 26, 32). The VasX *V. cholerae* effector encoded in the Aux 2 cluster is a large protein that contains a C-terminal colicin domain effective at eliminating both bacterial and eukaryotic cells (19, 20). Since it is predicted to form large pores, VasX permeabilizes cells and allows passage of molecules like PI into the cell (19). The *Pseudomonas aeruginosa* Tse4 and the *Serratia marcecens* Ssp6 effectors are both relatively small proteins that form ion-selective pores but do not allow larger molecules like PI to enter cells (53, 54). Recently, *Vibrio parahaemolyticus* has also been shown to harbor T6SS effectors that disrupt cellular membranes (58).

Importantly, the *V. cholerae* TpeV T6SS effector we describe in this study does not contain predicted domains or motifs with known functions and its sequence does not share homology to any previously described T6SS effectors. We provide evidence that TpeV is a T6SS toxin that can be used by *V. cholerae* cells to permeabilize target cells and disrupt the cell membrane potential (Fig. 2, Fig. 3). The cell membrane potential is essential for ATP synthesis, cell division and membrane transport (59–61). Therefore, TpeV-mediated toxicity is likely to inflict substantial damage to target cells by perturbing multiple essential processes.

We hypothesize that TpeV could permeabilize cells by forming pores. Pore-forming toxins (PFTs) are widespread among all kingdoms of life (62–66). Based on the secondary structure of the membrane spanning domain, two major classes of pore-forming toxins (PFTs) have been described: α-PFTs and β-PFTs (62, 64, 67). α-PTFs include the *E. coli* colicin and cytolysin A families, while β-PFTs are found in many Gram-positive bacterial species and contribute to the virulence of pathogens like *Staphylococcus aureus* and *Clostridium perfringens* (64, 66–68). Our homology predictions suggest that TpeV might harbor a peptidoglycan-binding OmpA-like domain (49). RmpM is a *Neisseria meningitidis* periplasmic protein that also possesses an OmpA-like domain (50). Experimental evidence indicates that RmpM stabilizes oligomeric porins in the outer membrane (50, 69). Rather than form new pores, it is also possible that TpeV might interact with and disrupt the normal functions of existing porins or channels in the membranes of target bacteria. Future experiments will determine if TpeV forms pores or employs other mechanisms that damage membranes and permeabilize cells.

In strain BGT49, the Aux 4 cluster and a restriction modification system are found near a prophage integrase and a transposase (Fig. 1A). This suggests that the genes are located on a mobile genetic element that can be transferred between bacterial cells to confer competitive advantages against phages and other bacterial cells (32). The *Vibrio cholerae* T6SS Aux 3 cluster was also recently shown to be located on a mobile genetic element (29). Our results showing that *V. cholerae* strain C6706* can use the Aux 4 cluster to kill parental cells supports the hypothesis that the Aux 4 cluster can be transferred to confer competitive advantages. This hypothesis is further supported by our observation that TpeV homologs are found close to other T6SS genes in many bacterial species, including human pathogens (*Providencia rettgeri, Proteus mirabilis, Citrobacter koseri, Yokenella regensburgei, Serratia fonticola, Salmonella enterica*, and *E. coli*), animal pathogens (*Vibrio anguillarum* and *Photorhabdus luminescens*) and plant pathogens (*Pectobacterium parmentieri*) (70–77) (Fig. 4). TpeV homologs found in other bacterial species are also located near transposase-like genes (data not shown).

All known T6SS toxic effectors are neutralized by cognate immunity proteins, which are generally encoded by genes adjacent to effectors (19, 78). We found that *tpiV*, the gene found immediately downstream of *tpeV*, confers immunity to target cells against *tpeV-*mediated toxicity (Fig. 1B and 2B). SignalP predicts that TpiV encodes a periplasmic Sec-tag, which is expected since TpeV exhibits its toxicity when delivered to the periplasm of target cells (Supp. Fig. 2, Fig. 3A). In other species, we observed that multiple putative immunity proteins can be found near TpeV homologs (Fig. 4). Additional studies are required to confirm which predicted TpiV-like proteins are the cognate immunity factors for the TpeV homologs.

In conclusion, we demonstrate that the T6SS Aux 4 cluster found in many *V. cholerae* isolates encodes a toxin that can be used to eliminate competitor bacteria. TpeV is a T6SS effector that permeabilizes target bacteria and disrupts the membrane potential, leading to severe cellular intoxication. However, target cells expressing TpiV are protected and resist TpeV-mediated toxicity. Finally, we find that TpeV homologs are widespread among Gram-negative bacteria, suggesting the protein represents a novel and potent antimicrobial agent of interest for further studies. Understanding the molecular mechanisms of antimicrobial toxins that drive competition and antagonism could lead to the development of novel biotechnology and medical applications.

## METHODS

### Bacterial strains and plasmids

Plasmids were constructed using standard molecular biology techniques. Gibson mix reagents, restriction enzymes and polymerase were used as recommended by manufacturers (Promega and New England Biolabs). Plasmids were verified by PCR and Sanger sequencing (Eurofins). *V. cholerae* C6706 mutant strains were made using pKAS allelic exchange methods, as described previously (79). *V. cholerae* strain BGT49 mutant strains were made using natural transformation, as described previously with modifications (80, 81). Briefly, overnight cultures were back-diluted in fresh LB medium for approximately 1 hour and then statically incubated overnight at 30°C in liquid LB medium with a sterile crab shell fragment. Crab shells were transferred and incubated in fresh LB medium containing 30-50 µg of a plasmid engineered to encode ∼1000bp flanking regions to replace the desired genes with an antibiotic cassette. Cells were incubated statically overnight at 30°C liquid LB medium and then spread on antibiotic plates to select for transformants. BGT49 mutants were confirmed by PCR and antibiotic resistance. Bacterial strains and plasmids used are listed in Supplementary Table 2.

### Bacterial Competition Assays

Bacterial cultures were grown overnight in liquid LB medium at 37°C with shaking. Overnight cultures were back-diluted and incubated in liquid LB medium at 37°C with shaking for 3 hours. Bacterial cultures were then normalized to an OD_600_ absorbance of 1. If strains harbored plasmids, cultures were grown overnight with antibiotics to maintain plasmids and 100 µM IPTG if plasmids contained an inducible promoter. If strains were grown in media containing antibiotics, liquid cultures were then washed three times with fresh LB medium before they were co-cultured. A 50 µL mixture aliquot of ratio of 10:1 killer:target cells was spotted on a 0.22 µm pore size filter paper, which was placed on LB agar media and incubated at 37°C. After 3 hours, filters were vortexed in sterile LB media for 30 seconds. 100µL of serial dilutions were then spread on plates containing the required antibiotic to select for target cells. Data from three co-cultures were used to determine significance. Results are representative of at least two independent experiments.

### Confocal microscopy

Overnight cultures were back-diluted 1:100 for 3 hours in liquid LB medium. Samples were then normalized to an OD_600_ of 10. 1 µL aliquot of 10:1 killer:target cell mixture was spotted on top of a dry 8-μL aliquot of propidium iodide (100 μg/mL) on an LB agar pad. Nikon A1R confocal microscope using a Perfect Focus System with a 40x objective (Plan Fluor ELWD 40x DIC M N1) was used to stabilize the focus in the plane of the colony growth. Cells were imaged at 96-100% humidity and 37°C. Images were processed using ImageJ. Results are representative of at least three independent experiments.

### Membrane permeabilization assays

Bacterial cultures of *E. coli* Shuffle T7 Express (New England Biolabs) cells carrying either a control plasmid or a periplasmic *tpeV* construct were grown overnight in liquid LB medium supplemented with 0.2% glucose and ampicillin at 37°C with shaking. Cells were washed three times with LB and 100x back-dilutions were made in fresh liquid LB medium with 500 µM IPTG and ampicillin. Strains were incubated at 37°C for 2 hours, washed three times with PBS and normalized to an OD_600_ of 1. 100µL of each culture was incubated with 1 µL propidium iodide (100 μg/mL) for 15-30 minutes. Fluorescence values were taken on a Synergy BioTek plate reader using an excitation λ = 535 nm and emission λ = 617 nm and normalized by subtracting the average values from samples with propidium iodide but no cells. Data represents the averages obtained from seven biological replicates from two independent experiments.

### Membrane potential assays

Bacterial cultures of *E. coli* Shuffle T7 Express (New England Biolabs) cells carrying either a control plasmid or a periplasmic *tpeV* construct were grown overnight with shaking at 37°C in liquid LB medium supplemented with 0.2% glucose and ampicillin. Cells were washed three times with LB and 100x back-dilutions were incubated at 37°C for 2 hours in fresh liquid LB medium with 500 µM IPTG and ampicillin. Cells were again washed three times with PBS and normalized to an OD_600_ of 1 in PBS. Cells were incubated for 30 minutes in the dark with DiBAC4(3) at a final concentration of 10 µM and washed with three times with PBS. Fluorescence values were taken on a Synergy BioTek plate reader using an excitation λ = 490 nm and emission λ = 516 nm. Data represents the averages obtained from six biological replicates from two independent experiments.

For co-culture measurements of membrane potentials, overnight cultures of the indicated *V. cholerae* BGT49 strains were back-diluted and incubated in liquid LB medium at 37°C with shaking for 3 hours. Bacterial cultures were then normalized to an OD600 absorbance of 1. A 50 µL mixture aliquot of ratio of 1:1 killer:target cells was spotted on a 0.22 µm pore size filter paper, which was placed on LB agar media and incubated at 37°C. After 3 hours, filters were vortexed in sterile LB media for 30 seconds. Cells were washed three times with PBS, normalized to an OD_600_ of 1, incubated for 30 minutes in the dark with DiBAC4(3) at a final concentration of 10 µM and washed with three times with PBS. Fluorescence values were taken on a Synergy BioTek plate reader using an excitation λ = 490 nm and emission λ = 516 nm. Data represents the averages obtained from six biological replicates from two independent experiments.

### Bioinformatic analyses

The HHMER server was used to search for homologs of TpeV in the UniProtKB database (82). Selected homologs were aligned using MUSCLE (83). A phylogenetic tree was constructed using PhyML with 100 bootstrap values and visualized using PRESTO (84–86). Putative immunity proteins were predicted based on homology to TpiV and genomic location. Truncated VgrG-like genes with stop codons were observed in some species but were excluded from Fig. 4.

## ACKNOWLEDGEMENTS

We thank Dr. Jacob Thomas, Dr. Mackenzie Martin, Dr. Athéna Patterson-Orazem, Dr. Dustin Huard and Tong Yu for advice, experimental help and useful discussions.

BKH would also like to thank funding from the Georgia Institute of Technology School of Biological Sciences, NSF (MCB 1149925 and BMAT-2003721) and BSF (2015103). GS would like to thank the German National Academy of Natural Sciences Leopoldina (LDPS 2017-03).

## SUPPLEMENTARY FILES LEGENDS

**Supplementary Figure 1. TpeV Phobius transmembrane helix prediction**.

**Supplementary Figure 2. TpeV SignalP prediction**.

**Supplementary Figure 3. TpiV SignalP prediction**.

**Supplementary Table 1. Complete list of all identified TpeV homologs using the PHMMER algorithm**.

**Supplementary Table 2. Strains and plasmids used in this study**.

